# Geology and climate drive alpine plant compositional variation among peaks in the Cascade Range of Washington

**DOI:** 10.1101/2024.12.20.629768

**Authors:** Erik W. Ertsgaard, Nicholas L. Gjording, Jonathan D. Bakker, Joseph A. Kleinkopf, David E. Giblin

## Abstract

Alpine areas are host to diverse plant communities that support ecosystems through structural and floral resources and persist through specialized adaptations to harsh high-elevation conditions. An ongoing question in these plant communities is whether composition is shaped by stochastic processes (e.g., dispersal limitations) or by deterministic processes (e.g., climate, geology), and if those processes select for common phylogenetic clades across space. This study evaluates the drivers of dissimilarity in alpine vascular plant communities across 32 peaks in the Cascade Mountain Range of Washington State and examines the effects of incorporating phylogenetic relatedness to these conclusions. We documented an average of 54 species per peak and used our overall inventory of 307 taxa to construct a phylogenetic tree for the entire mountain range plant community sampled. We used multivariate techniques to quantify the phylogenetic and taxonomic differences between alpine plant communities and to relate those differences to each peak’s climate, geology, and topography. Our models indicate that the age of each peak’s parent material formation, precipitation, latitude, and temperature had the largest role in shaping alpine plant communities relative to the baseline effects of distance between peaks and time of sampling. A unique result was a distinct plant community in peaks with ultramafic geologic parent material formed in the Paleozoic Era, which has an extreme geochemistry that we found to form evolutionarily distinct lineages compared to all other peaks. With changing climate conditions and disturbance regimes, understanding facets of alpine plant communities like species turnover, geologic endemism, and responses to precipitation changes are vital to conserving these ecosystems.

## Introduction

Alpine plant communities contribute a disproportionately high amount of global vascular plant diversity and exist at the elevational limits of plant distributions [1,2]. In turn, the structure and resources provided by alpine plants in these extreme environments support diverse communities of pollinators, herbivores, and fungi at their elevational limits. Though alpine ecosystems are relatively protected from human development, climate change and recreation pose major threats to their diversity. Trends of plants moving to higher latitudes and elevations in accordance with climate shifts have been widely speculated and demonstrated by plant biogeographers [3–5], yet alpine systems have been theorized to contain adequate topographic heterogeneity [6] and/or plant trait diversity [7] to buffer these effects or decouple this direct climate-community interaction [8].

A useful way to explore patterns in mountain range plant communities is the study of beta-diversity: spatial variation in community composition. Such studies have been applied to alpine plant communities to investigate patterns in climate to determine whether neutral processes like dispersal limitations drive composition more than broad environmental filters like climate [6,9–12]. Dispersal limitation is of special concern for alpine systems as many theorize their isolation in matrices of other habitats create conditions similar to those of islands [13–15]. Smithers and Oldfather et al. [16] examined beta-diversity across climatic gradients at multiple scales in the White Mountains of California and found strong elevation gradients among peaks in both community dissimilarity and species’ climatic tolerances. However within peaks these relationships were much weaker. Malanson et al. [17] attempted to decipher whether neutral or deterministic patterns of plant composition occurred across the Rocky Mountains by contrasting multilevel (hierarchical) and multiscale (scales separated) models, finding confounding results between broad-scale climate and microclimates that point to a need for further study.

Environmental determinants of plant distributions go beyond climate. Distance is commonly used to relativize climate findings, identify dispersal limitations, or test the presence of broad vegetation patterns [10,11,17,18]. Geology has been identified as an important driver of composition, but because it does not conform to a gradient, it appears to contribute to compositional stochasticity in models that exclusively investigate climate [5,19]. In a continent- wide exploration of species turnover, Malanson et al. [19] found the strongest correlation between distance and turnover in the Rocky Mountains and much weaker relationships in other ranges like the Cascades. This was hypothesized to be a result of the relative youth of the Cascade Range yielding greater geologic diversity and subsequent stochasticity. Few studies have examined the strength of environmental (climate) gradients in relation to distance, geology, and topography to holistically resolve the determinants of alpine plant community composition. And those few studies span a small diversity of mountain ranges, limited to the North American Rockies [9,11,20], European Alps [13,21], and Himalayan Plateau [12].

Another way to improve our understanding of how alpine plant communities are structured is to consider different measures of compositional dissimilarity. Conventional taxonomic measures treat each taxon equally, but phylogenetic distances incorporate phylogenetic relatedness and can provide insight into questions of community formation, trait evolution, and niche structure [22]. Comparisons of these measures can address the question of whether neutral or deterministic processes shape alpine systems as predicted by island biogeography [9,13,23,24] but yield mixed findings. Jin et al. [9] found phylogenetic turnover was consistently lower than taxonomic (species-equal) turnover across Rocky Mountain alpine plant communities, and that environmental variables were more predictive than spatial variables, suggesting significant niche conservatism in these systems. This is supported by other findings of phylogenetic clustering in alpine systems [13,23]. However, studies have simultaneously failed to reject neutral processes at the mountain-range level, indicating either complexity in environmental determinants that were not tested or true stochasticity among summit plant communities [13,24]. Yu et al. [12] found phylogenetic beta-diversity of alpine plants of the Qinghai-Tibet Plateau to be linked to distance and climate, but did not include geologic variables and had a relatively coarse spatial resolution. Further examination of phylogenetic patterning of alpine plants with breadth in potential environmental determinants and across varying mountain ranges is needed to understand the evolutionary and contemporary processes driving community composition.

We examined compositional variation among alpine plant communities to address three specific objectives:

1. Quantify the degree of relation between alpine plant communities and the importance of climatic, geologic, and topographic variables to dissimilarity between plant communities while controlling for distance between peaks, thus obtaining a holistic understanding of the environmental filters on plants occurring in the Cascade Range alpine zone.
2. Compare results when applied to taxonomic and phylogenetic dissimilarities to reveal the importance of processes such as phylogenetic clustering.
3. Identify plant species indicative of distinct alpine communities in the Cascade Range.

## Methods

### Ethics Statement

All surveys were conducted on U.S. Forest Service lands, which are public lands, under the terms of permits FS-2400-008 and FS-2700-4 (OWF009). No protected species were sampled. This field research was approved by the University of Washington. As a field-based study of vascular plants, this research was not subject to an internal review board at the University of Washington.

### Study Area

The Cascade Range is a relatively young formation of volcanic peaks on the West Coast of North America, stretching from southern British Columbia to northern California. The range contains several volcanoes of notable prominence (e.g., Mt. Rainier, 4392 m), but is composed mostly of peaks between 1200 and 2700 meters in elevation. Due to their orientation perpendicular to the prevailing winds, the Cascades experience a strong orographic effect with much higher precipitation on the windward (western) than the leeward (eastern) side of the Cascade crest [25]. Notable geologic diversity is stored within the range due to volcanic events of different ages and to series of terranes of varying lithology [26].

### Floristic Surveys

For this study we define the alpine zone as that area on each peak above the continuous treeline. We conducted floristic surveys in the alpine zone of 32 peaks within the Washington portion of the Cascade Range (Fig 1). Sampled peaks were selected on the basis of multiple criteria: 1) no technical gear (e.g., ropes) or technical ascents (e.g., glacier travel) required to reach the summit; 2) representation of east and west sides of the Cascade Range crest to capture variation in precipitation; and 3) representation along north-south axis of Cascade Range in Washington to capture geologic and topographic variation. Floristic surveys were conducted between July 1st and September 1st in 2021-2023, and peaks ranged between 1757-2512 m elevation.

**Fig 1.**
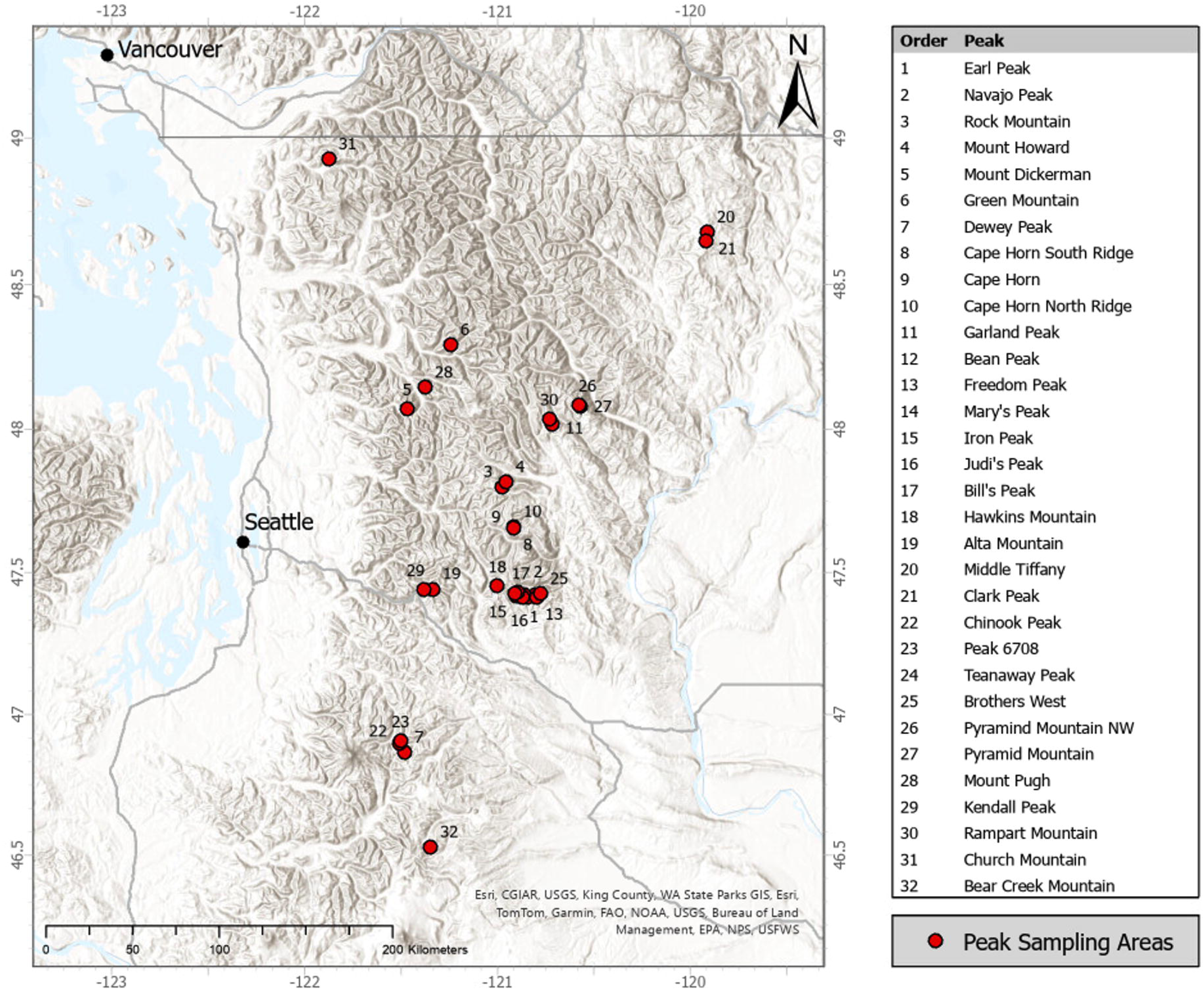
Peaks sampled. Locations of sampled peaks across the Cascade Mountain Range of Washington State.

In the Cascade Range in Washington where we sampled, treeline depends on aspect and precipitation. Consequently, a higher peak does not necessarily mean a larger alpine area. On each peak we thoroughly searched all accessible terrain recording taxa until further searching generated no new taxa. Inaccessible terrain typically comprised cliffs and sheer faces with little or no vegetation. Alpine areas on sampled peaks were relatively small in area, ranging in size from 372 m^2^ to 43717 m^2^, such that an entire peak could be sampled in a single outing.

Each taxon encountered was identified in the field by sight where possible and added to the peak list. Voucher specimens were made for all taxa not easily identified in the field (e.g., Poaceae, Cyperaceae, Brassicaceae), and taxa representing novel distribution records. No protected species were sampled. Those specimens were later identified at the University of Washington Herbarium in Seattle, WA using the contemporary regional flora manual [27]. With herbarium specimens available for comparison and experts in the regional flora on the field teams, taxa were identified with unique accuracy both in the field and in the herbarium. For each taxon, qualitative information on abundance, phenology, and aspect was recorded. For each peak, geocoordinates, elevation, sampling date, sampling area, dominant habitat, and dominant species were also recorded.

Pacific Northwest summers are relatively dry, with snowmelt providing the majority of soil moisture for alpine plant growth and reproduction. Consequently, flowering begins shortly after snowmelt. Our sampling occurred over a large time window because elevation differences

determine timing of snowpack melting. Lower elevation peaks are snow-free in June, and flowering begins shortly after snowmelt. Moreover, the greatest risk of overlooking taxa during our surveys was sampling too early, and taxa in several families (e.g., Brassicaceae, Cyperaceae) cannot be accurately identified in flower because fruits are required for accurate identification.

### Explanatory Variables

Climate, geology, and topography data were collected for each peak. Variables were chosen to represent the holistic suite of landscape-level processes that could influence plant communities. Because these analyses were not initially in the scope of our floristic surveys, most variables were adapted from public datasets. Definitions of all variables and their sources are described in full in S1 Table.

For our analyses, space and time variables were included as covariates; these include latitude, longitude, sampling area, year, and date of sampling. While spatial variables can address ecological questions like the presence of dispersal limitations, both space and time covariates normalize latter evaluations of environmental variables that may be conflated by these effects.

Latitude, longitude, and elevation approximate in the influence distance between peaks, but each can also contain other relevant gradients (e.g., east-west orographic precipitation patterns).

Sampling area addresses issues of size differences among alpine areas and was estimated from GPS tracks of crew members as shown in Fig 2. Because our sampling effort occurred over a single day, the date of sampling could affect which plants were present and identifiable within a year, as well as the crew doing the identifications between years. To incapsulate both unintended influences of our sampling design, we divided the date of sampling into the year (2021, 2022, or 2023) and the number of days into the field season (after the earliest date of all three years, June 29th).

**Fig 2.**
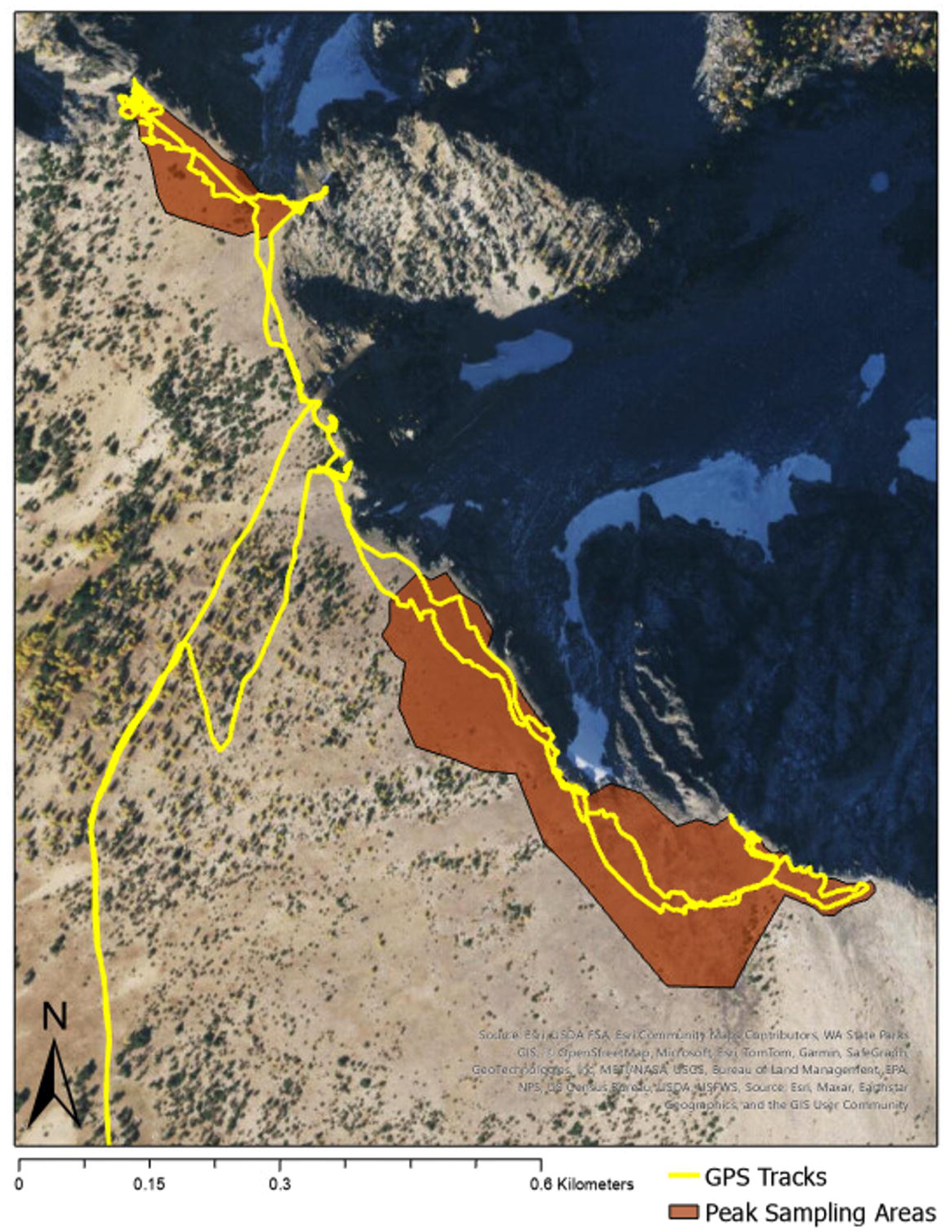
Peak survey method. GPS tracks of a field crew member’s sampling of two peak alpine areas (top left is Pyramid Peak Northwest and bottom right is Pyramid Peak) depicting the sampling design of identifying plants in all accessible areas above the continuous treeline. Peak sampling area polygons were drawn from GPS tracks, satellite imagery, and recollection of the entire field crew’s sampling extent and were used to generate topographic and covariate variables. Satellite image provided by USGS/The National Map (public domain).

Climate variables attempt to represent the broad variability in alpine weather patterns and climatic processes while avoiding multicollinearity. We followed the selection criteria of Bakker et al. [27], who subset the 19 WorldClim bioclimatic variables (https://www.worldclim.org/data/worldclim21.html) to only include the six least correlated predictors for grasslands across the world: mean annual temperature (MAT), mean annual precipitation (MAP), annual temperature range, temperature of the wettest quarter, temperature seasonality (standard deviation of monthly temperature values), and precipitation seasonality (Coefficient of variation of monthly precipitation values). These are suitable for our analyses because they cover both the intensity and variability of plant-relevant climate patterns while being previously modelled across our study area at a sufficient resolution. Each variable was calculated using monthly temperature and precipitation data from 30-year normals (1991-2020) modeled by the PRISM Climate Group with 800 m resolution [28].

Geologic variables were drawn from the Washington Department of Natural Resources Geologic Information Portal (https://www.dnr.wa.gov/geologyportal), with the bedrock parent (material intrusive, volcanic, heterogenous-metamorphic, or other outlier) and the age of its formation (Paleozoic, Mesozoic, or Cenozoic) coded categorically [29]. Being the two components of regional geologic maps, parent material and its age define the coarse-scale patterns of geology most accessible to managers or citizens. We also include soil development as a representation of fine-scale differences in edaphic habitat (e.g., active scree vs. stable tundra soil).

Topographic variables sought to quantify the potential effects of microclimates or other resource complexities on broader biogeographic patterns. We drew from 10-meter digital elevation models using the USGS TNM Downloader (https://apps.nationalmap.gov/downloader/) to quantify the elevation range and topographic complexity of each peak. The Rumple Index from Kane et al. [30] was adopted to quantify surface complexity of the peak: the ratio of three-dimensional area to two-dimensional surface area. While previously used to measure the complexity of tree canopies, Rumple Index fits our intention to measure the roughness or irregularity of a peak’s three-dimensional shape that correlate to the amount of microclimates.

### Phylogenetic Data

We constructed a phylogenetic tree of all recorded taxa using the *V.Phylomaker2* package [31], drawing from the GBOTB.extended.WP.tre mega-tree [32] with nomenclature standardized with the World Plants (WP) Database (https://www.worldplants.de). Mega-trees are large phylogenetic trees based on molecular sequence data available in public databases. Although this package does not track the most up-to-date APG IV nomenclature, the phylogeny is resolved at the genus level and thus is comparable to other community ecology studies [33]. To adapt the mega-tree to only include taxa we observed, the `phylo.maker` function “pruned” the tree to taxa names matching those in the mega-tree and “binded” taxa to corresponding species or genera if the (infra)species is not found in the mega-tree. As such, the phylogenetic difference between two taxa in this tree is the number of branches removed they are from one another within the alpine species pool (Fig 3).

**Fig 3.**
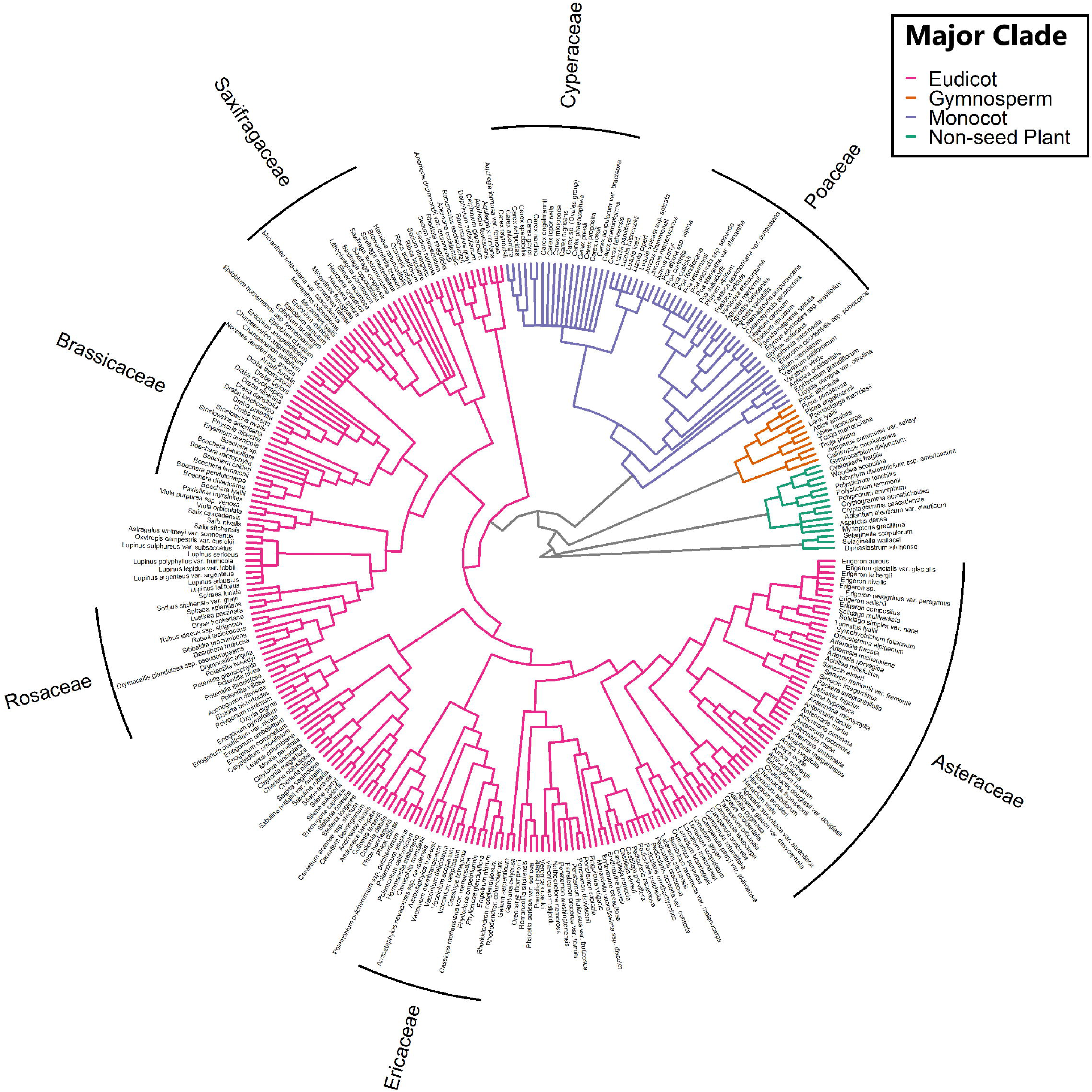
Phylogeny of species sampled. Phylogenetic tree of all 307 taxa recorded in this study, compiled from a mega-tree and used to calculate the Unifrac distance between peaks.

### Statistical Analyses

All analyses were performed using R statistical software [34]; the full reproducible code is available in Supplementary Materials. Vascular plant community dissimilarity between peaks was quantified using Jaccard dissimilarity and UniFrac distance. Jaccard dissimilarity is appropriate for presence-absence data and compares any two peaks in terms of the proportion of taxa that were present on only one of the peaks. This dissimilarity was calculated using the `vegdist` function within the *vegan* package [35].

UniFrac distance is commonly used for testing phylogenetic relationships in microbial communities [36], but has also been utilized for vascular plant communities [13,24] as a measure of terminal beta diversity. Consistent with the methods outlined in Cadotte and Davies [37], we computed unweighted dissimilarity (without abundance information), incorporating the fraction of total unshared branch lengths derived from our phylogenetic tree using the *GUnifrac* package in R [38]. The resulting phylogenetic distance matrix contrasts from the prior taxonomic distances in the incorporation of phylogenetic relatedness in taxa not shared between peaks. That is, taxa are not singularly matched via their names but are evaluated in the context of shared evolved traits and genes within clades. In other words, taxa of the same genus or family are more morphologically similar than those in different phyla and therefore may react similarly to environmental conditions. These phylogenetic distance results can then be compared to taxonomic results to see if biogeographic patterns of community composition stem from evolved traits (i.e., niche conservatism).

Each measure of compositional dissimilarity was evaluated against geographic distance in a Mantel test with Pearson’s product-moment correlations and 100,000 permutations as coded in the `mantel` function of the *vegan* R package [35]. The resulting correlations had nonzero slopes and were significant, so spatial autocorrelation terms—latitude and longitude—were included in subsequent analyses.

To test the significance of our explanatory variables on the composition of alpine vascular plant communities, we conducted a series of permutational multivariate analysis of variance (PERMANOVA) models for each distance measures. PERMANOVA permutes variable identities to assess the significance of partitions in multivariate response data that correspond to categorical and continuous predictors [39]. Here, continuous variables were normalized to express them on the same scale. For our analyses, we employed a top-down model selection framework that evaluated all possible configurations of predictor terms by their Aikaike information criterion, corrected for sample size (AIC_C_), using the *AICcPermanova* Package [40].

We conducted all model testing with *vegan*’s `adonis2` function [35], each using 1,000 permutations and Type III Sums of Squares to test the marginal effect of each term above all others. Final models included all with AIC_C_ values within two units of the top-performing model [41] and excluded those with high multicollinearity (VIF>5). To assess the significant trends across all final models, we utilized model-averaging to generate AIC_C_-adjusted partial R^2^ values that represent the average explanatory power of each term compared to when it is not included.

We conducted non-metric multidimensional scaling (NMDS) ordinations of the alpine plant community to visualize significant relationships as identified in the PERMANOVA analyses for each distance measure. The dimensionality of our final solutions was selected by iterating between one and five dimensions to select the value which balances the plot’s interpretability while maintaining a stress value from which we can draw meaningful conclusions. We chose to use three dimensions in our final solutions and to plot the first two dimensions of these solutions. The NMDS parameters met or exceeded the minimum recommended values proposed by McCune and Grace [42] to ensure a global solution is met: 400 iterations, 40 minimum tries, and 100 maximum tries. Significant variables from the PERMANOVA analyses were overlaid onto the NMDS point cloud to illustrate compositional patterns among peaks.

To identify taxa associated with compositional distinct communities, we performed a simplified indicator species analysis (ISA) and threshold indicator taxa analysis (TITAN).

Simplified ISA compares the presence of taxa in a group compared to its presence in other groups; a strong indicator is present only in one group, not any others. Here, we use the method proposed by De Caceres et al. [43] that evaluates groups individually and together (e.g., Indicating Group A or Groups A and B). Significance of indicators is assessed through Monte-Carlo randomized permutations [44]. The model results include an Indicator Value, the product of the species’ relative specificity and fidelity to a group, and a permuted significance value.

Although Simplified ISA is adapted from a classical ISA that incorporates species’ abundance values to only require presence-absence information, Bakker et al. [45] found this resulted in minimal difference in results. Within the `multipatt` function of the *indicspecies* R package [46], we used 9999 permutations and considered species with IV>0.5 as strong indicators at α = 0.05. Because our sampling design was non-hierarchical, we did not restrict permutations.

TITAN identifies indicator taxa for continuous variables under a framework similar to a regression tree: iteratively splitting environmental variables into two groups to identify species that increase or decrease as the environmental variable changes. TITAN uses bootstrap permutation testing and generates values associated with the species purity (consistency of directionality) and reliability (frequency of significance in bootstrap permutations). We used the function, `titan`, within the *TITAN2* R package [47]. To decrease variation among runs, we used 1000 permutations and 1000 replicates in bootstrap resampling. We used a cutoff value of 0.95 for both purity and reliability values consistent with α = 0.05.

## Results

Across 32 peaks in the Cascade Range, we documented 307 vascular plant taxa (Fig 3).

Taxonomic distance, measured by Jaccard Dissimilarity, was consistently lower than phylogenetic distance, measured by Unifrac Dissimilarity, while retaining overall trends in between-peak dissimilarity (Fig 4).

**Fig 4.**
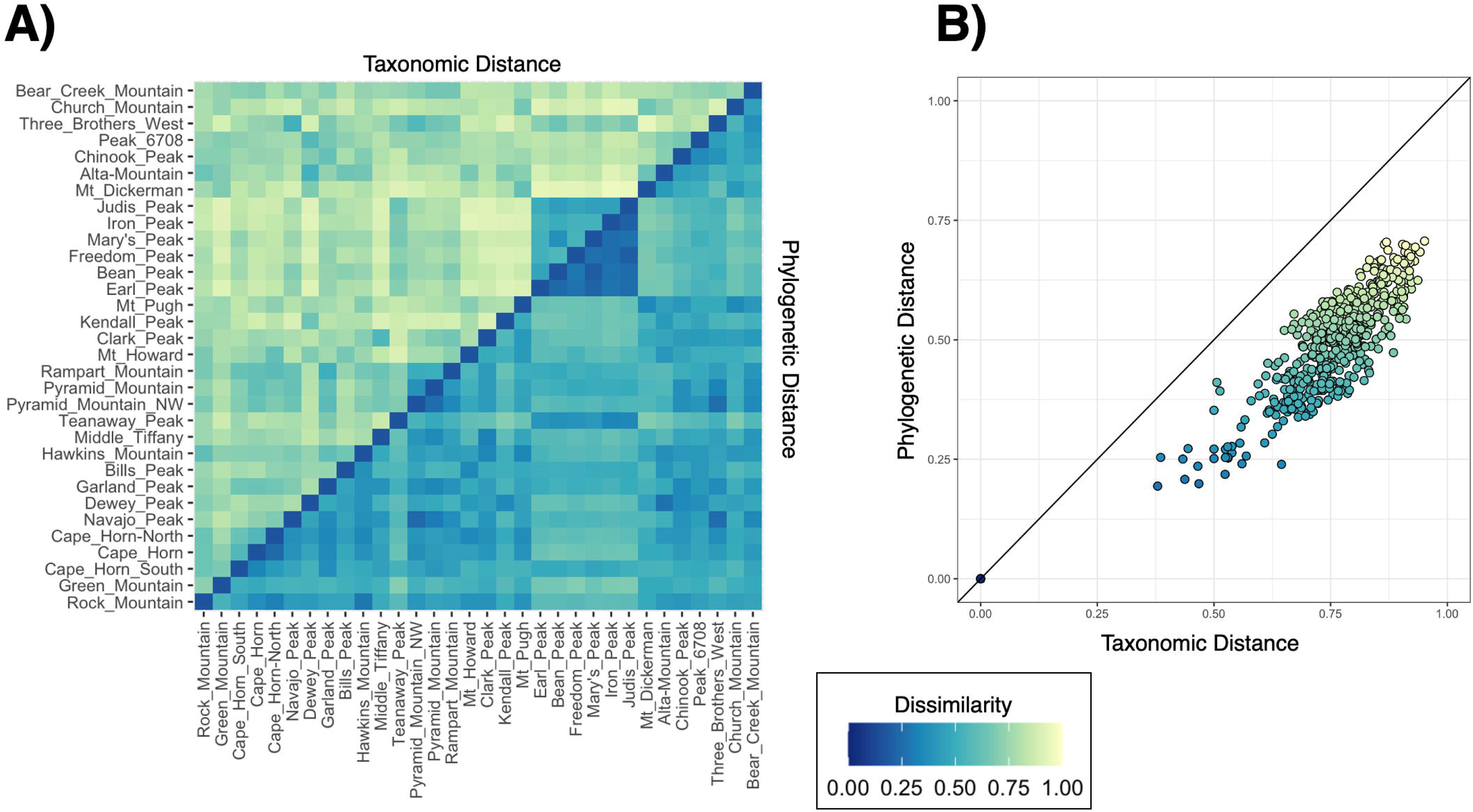
Jaccard and Unifrac distances between peaks. (A) Heatmap illustrating the relationship between Jaccard and Unifrac distances between peaks. Each tile is colored corresponding to the dissimilarity value between its two peaks, with Jaccard Dissimilarity values in the top-left half and Unifrac Dissimilarity in the bottom-right half. Peaks are ordered by geology to visualize its effects on dissimilarity for both distance measures. (B) Jaccard and Unifrac distances plotted together in a scatter plot with points colored by Jaccard dissimilarity values, and a 1:1 line to demonstrate the skewness toward taxonomic distance while maintaining a linear fit.

Mantel tests found significant spatial autocorrelation in both taxonomic (Mantel r =0.392, p <0.0001) and phylogenetic (Mantel r=0.174, p=0.0149) distance measures (Table 1). Our PERMANOVA analyses and systematic model selection yielded six variables which consistently correlate to alpine plant community composition across 184 final taxonomic models and 58 final phylogenetic models (Table 2). Period, or geologic age of parent material formation, explained the highest amount of variance in our taxonomic distance data (partial R^2^=0.087), while MAP, Latitude, Temperature of the Wettest Quarter, MAT, and Rumple Index explained much lower fractions in descending order. These variables remained consistent across phylogenetic distances, with the only notable difference between the two response datum being a much stronger effect of Period (partial R^2^=0.180). NMDS plots reflected these mirroring relationships in taxonomic and phylogenetic distances with significant explanatory variables overlaid, except for minor translocations of peaks in the data cloud (Fig 5). Convex hulls of geologic parent material levels illustrated tight (with the exception of one point) in peaks with parent material formed during the Paleozoic Era, and clear separation between all ages of formation (Figs 5A and 5C). MAP also demonstrated a clear gradient from high-precipitation peaks to low-precipitation peaks within NMDS point clouds (Figs 5B and 5D).

**Fig 5.**
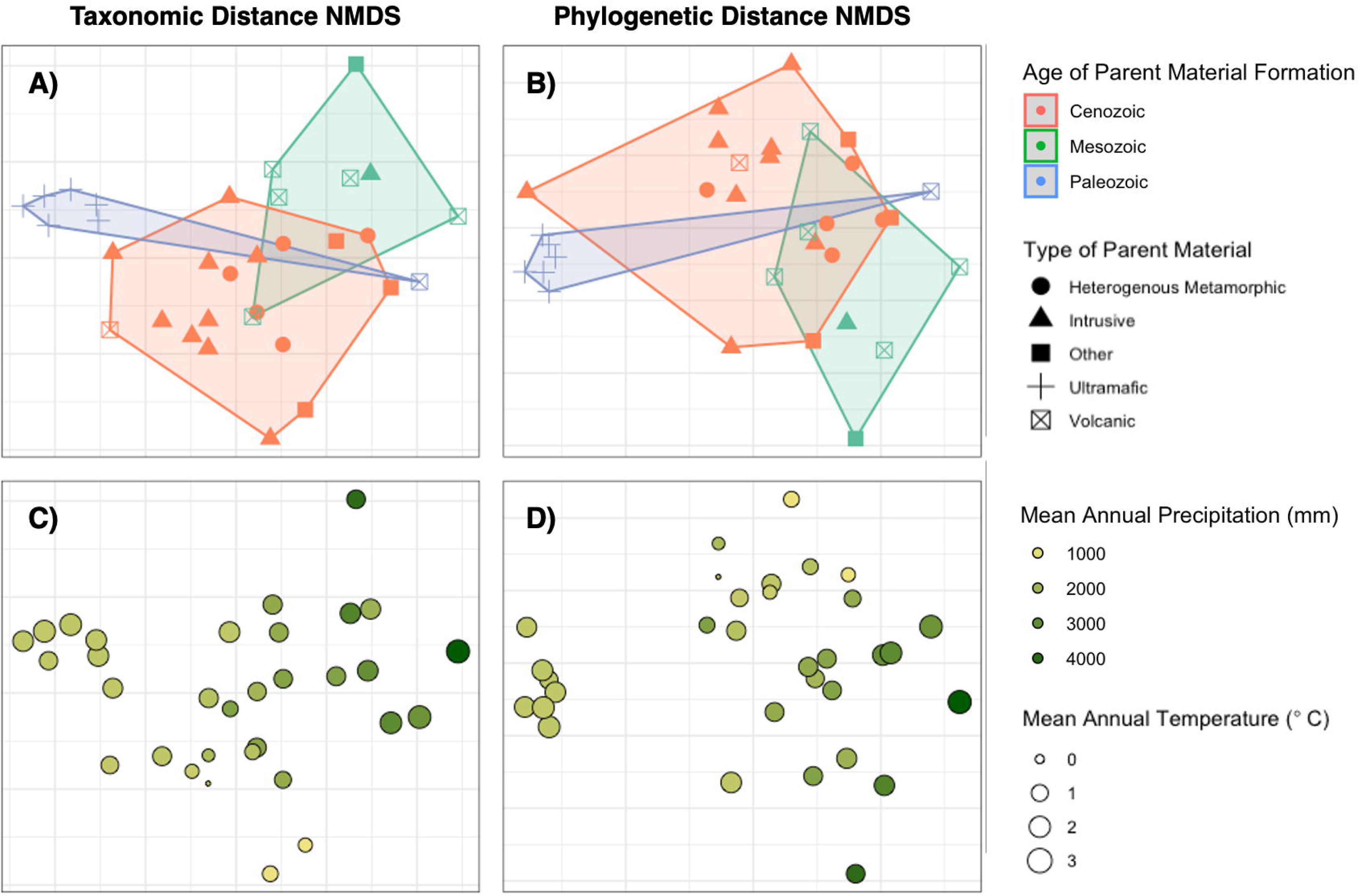
NMDS ordinations. Plots of NMDS ordinations for taxonomic distance (A and C) and phylogenetic distance (B and D) between alpine plant communities, with significant explanatory variables overlaid. Our NMDS of taxonomic distance and phylogenetic distance measures had stress values of 0.107 and 0.096 respectively. Age of Parent Material Formation groups are symbolized by point color, and convex hulls are drawn to encompass the entire group. Mean Annual Precipitation (MAP) values are symbolized by graduated colors, lighter yellows representing peaks with the most precipitation variability.

**Table 1.**
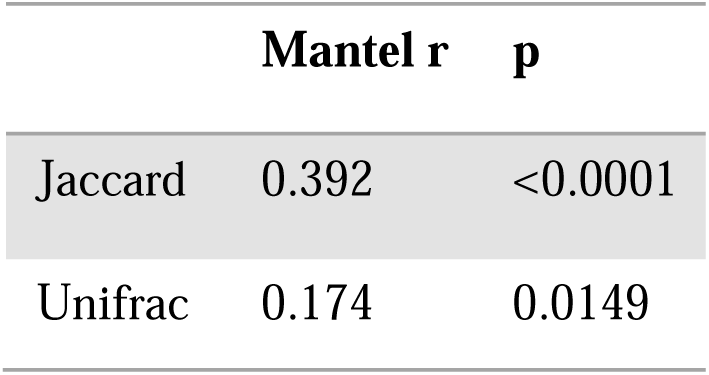
Mantel tests. Model results from Mantel tests of taxonomic (Jaccard) and phylogenetic (Unifrac) distances against geographic distance between peaks, using 100,000 permutations.

|  | Mantel r | p |
| --- | --- | --- |
| Jaccard | 0.392 | <0.0001 |
| Unifrac | 0.174 | 0.0149 |

**Table 2.** PERMANOVA analyses. . AICc-Adjusted Partial R^2^ values for final variables most predictive of alpine plant community composition, calculated from model averaging of final PERMANOVA models (ΔAICc<2 and VIF<5) for taxonomic (Jaccard) distances (184 models) and phylogenetic distances (58 models) as response matrices.

| Variable Name | Taxonomic (Jaccard) | Phylogenetic (UniFrac) |
| --- | --- | --- |
| | AICc-Adjusted Partial $R^2$ | AICc-Adjusted Partial $R^2$ |
| Period | 0.0872 | 0.1796 |
| **Mean Annual Precipitation (MAP)** | 0.0245 | 0.0225 |
| **Latitude** | 0.0168 | 0.0296 |
| **Temperature of Wettest Quarter** | 0.0149 | 0.0078 |
| **Mean Annual Temperature (MAT)** | 0.0094 | 0.0035 |
| **Rumple Index** | 0.0008 | 0.0023 |

ISA identified 36 taxa that are linked to specific parent material age groups in the Cascade Range in Washington that we sampled (Table 3). Seventeen taxa are Paleozoic peaks and another nine are notably intolerant of ultramafic peaks. TITAN identified 23 taxa associated with the continuum of MAP, 14 of which with increasing precipitation and nine of which with decreasing precipitation (Table 4).

**Table 3.** Relationship of taxa sampled to. parent material age. Taxa significantly linked to geologic period in our ISA. Taxa were filtered on indicator values (√IV > 0.707) and significance (p < 0.05).

| **Group** | **Taxa** | **$\sqrt{IV}$** | **p** |
| --- | --- | --- | --- |
| **Paleozoic** | *Aspidotis densa* | 0.926 | 0.005 |
|  | *Noccaea fendleri* ssp. *glauca* | 0.926 | 0.005 |
|  | *Polystichum lemmonii* | 0.926 | 0.005 |
|  | *Allium crenulatum* | 0.897 | 0.005 |
|  | *Claytonia megarhiza* | 0.871 | 0.005 |
|  | *Viola purpurea* ssp. *venosa* | 0.847 | 0.005 |
|  | *Oreocarya thompsonii* | 0.845 | 0.005 |
|  | *Androsace nivalis* | 0.825 | 0.005 |
|  | *Sabulina nuttallii* var. *nuttallii* | 0.794 | 0.005 |
|  | *Packera streptanthifolia* | 0.768 | 0.015 |
|  | *Pseudotsuga menziesii* | 0.761 | 0.005 |
|  | *Gallium serpenticum* | 0.756 | 0.005 |
|  | *Poa curtifolia* | 0.756 | 0.005 |
|  | *Eriophyllum lanatum* | 0.756 | 0.010 |
|  | *Erysimum arenicola* | 0.727 | 0.040 |
|  | *Boechera divaricarpa* | 0.722 | 0.005 |
|  | *Lomatium cuspidatum* | 0.722 | 0.010 |
| **Mesozoic** | *Eriogonum ovalifolium* var. *nivale* | 0.719 | 0.050 |
| **Cenozoic** | *Lomatium martindalei* | 0.893 | 0.005 |
|  | *Tsuga mertensiana* | 0.856 | 0.005 |
|  | *Vaccinium deliciosum* | 0.814 | 0.005 |
|  | *Luetkea pectinata* | 0.780 | 0.010 |
|  | *Luina hypolueca* | 0.748 | 0.005 |
|  | *Sedum divergens* | 0.742 | 0.015 |
| **Not Cenozoic** | *Castilleja elmeri* | 0.872 | 0.020 |
| **Not Mesozoic** | *Cryptogramma acrostichoides* | 0.817 | 0.010 |
|  | *Solidago multiradiata* | 0.716 | 0.050 |
| **Not Paleozoic** | *Trisetum spicatum* | 0.882 | 0.005 |
|  | *Festuca saximontana* var. *purpusiana* | 0.878 | 0.005 |
|  | *Eremogone capilaris* | 0.872 | 0.010 |
|  | *Juncus parryi* | 0.872 | 0.005 |
|  | *Penstemon procerus* var. *tolmiei* | 0.813 | 0.045 |
|  | *Tonestus lyallii* | 0.775 | 0.035 |
|  | *Phyllodoce glanduliflora* | 0.721 | 0.045 |
|  | *Selaginella scopulorum* | 0.721 | 0.045 |
|  | *Veronica cusickii* | 0.721 | 0.045 |
| **Paleozoic** | *Aspidotis densa* | 0.926 | 0.005 |
|  | *Noccaea fendleri* ssp. *glauca* | 0.926 | 0.005 |
|  | *Polystichum lemmonii* | 0.926 | 0.005 |
|  | *Allium crenulatum* | 0.897 | 0.005 |
|  | *Claytonia megarhiza* | 0.871 | 0.005 |
|  | *Viola purpurea* ssp. *venosa* | 0.847 | 0.005 |
|  | *Oreocarya thompsonii* | 0.845 | 0.005 |
|  | *Androsace nivalis* | 0.825 | 0.005 |
|  | *Sabulina nuttallii* var. *nuttallii* | 0.794 | 0.005 |
|  | *Packera streptanthifolia* | 0.768 | 0.015 |
|  | *Pseudotsuga menziesii* | 0.761 | 0.005 |
|  | *Gallium serpenticum* | 0.756 | 0.005 |
|  | *Poa curtifolia* | 0.756 | 0.005 |
|  | *Eriophyllum lanatum* | 0.756 | 0.010 |
|  | *Erysimum arenicola* | 0.727 | 0.040 |
|  | *Boechera divaricarpa* | 0.722 | 0.005 |
|  | *Lomatium cuspidatum* | 0.722 | 0.010 |
| **Mesozoic** | *Eriogonum ovalifolium* var. *nivale* | 0.719 | 0.050 |
| **Cenozoic** | *Lomatium martindalei* | 0.893 | 0.005 |
|  | *Tsuga mertensiana* | 0.856 | 0.005 |
|  | *Vaccinium deliciosum* | 0.814 | 0.005 |
|  | *Luetkea pectinata* | 0.780 | 0.010 |
|  | *Luina hypolueca* | 0.748 | 0.005 |
|  | *Sedum divergens* | 0.742 | 0.015 |
| **Not Cenozoic** | *Castilleja elmeri* | 0.872 | 0.020 |
| **Not Mesozoic** | *Cryptogramma acrostichoides* | 0.817 | 0.010 |
|  | *Solidago multiradiata* | 0.716 | 0.050 |
| **Not Paleozoic** | *Trisetum spicatum* | 0.882 | 0.005 |
|  | *Festuca saximontana* var. *purpusiana* | 0.878 | 0.005 |
|  | *Eremogone capilaris* | 0.872 | 0.010 |
|  | *Juncus parryi* | 0.872 | 0.005 |
|  | *Penstemon procerus* var. *tolmiei* | 0.813 | 0.045 |
|  | *Tonestus lyallii* | 0.775 | 0.035 |
|  | *Phyllodoce glanduliflora* | 0.721 | 0.045 |
|  | *Selaginella scopulorum* | 0.721 | 0.045 |
|  | *Veronica cusickii* | 0.721 | 0.045 |

**Table 4.**
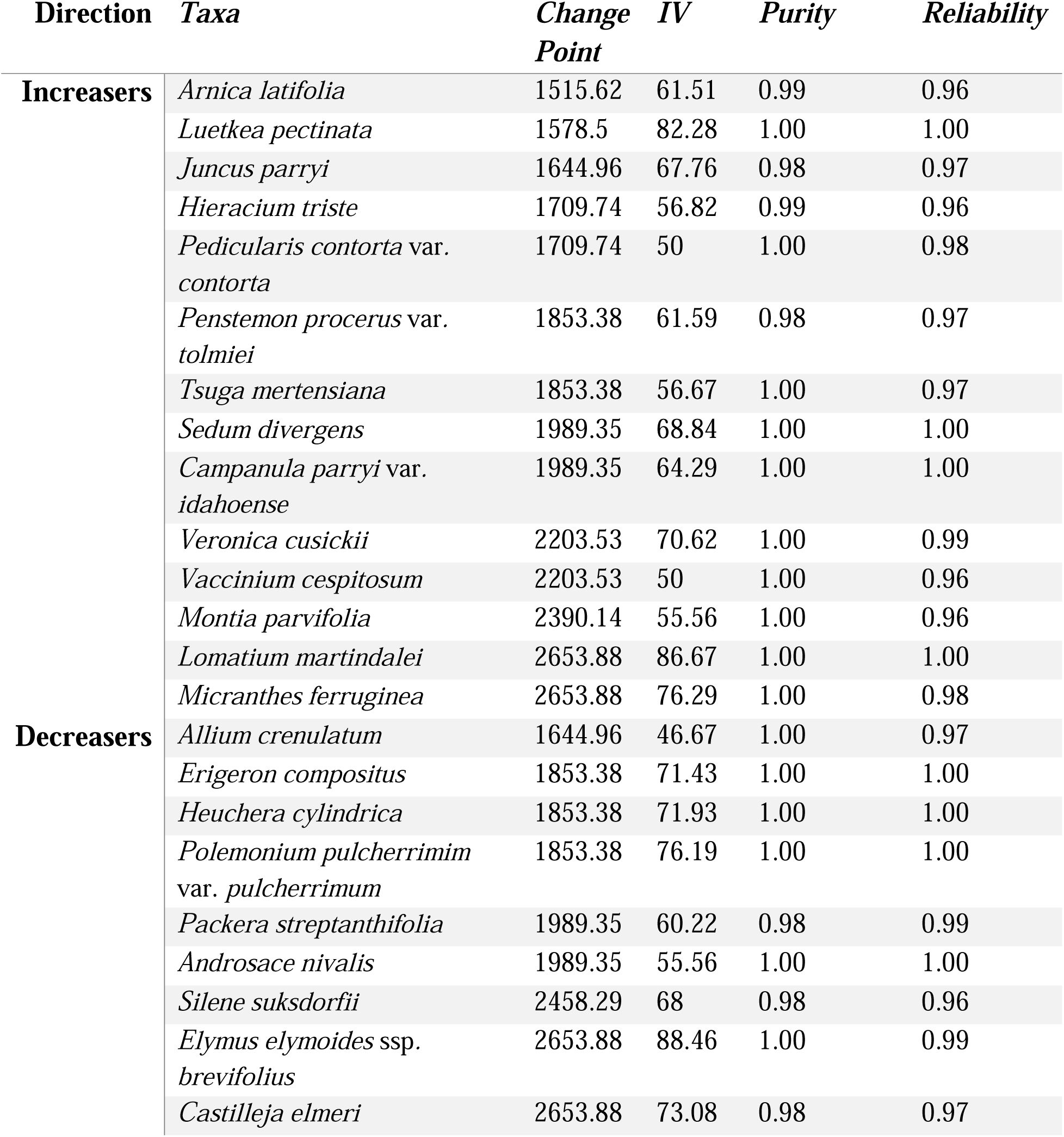
TITAN analyses.Taxa significantly associated with increasing or decreasing MAP in our TITAN. Taxa were filtered based on indicator values (IV) above 45 and purity and reliability values above 0.95. Change point values correspond to the MAP value (in mm) a species is more likely to be on the increasing or decreasing side of.

| Direction | Taxa | Change Point | IV | Purity | Reliability |
| --- | --- | --- | --- | --- | --- |
| **Increasers** | *Arnica latifolia* | 1515.62 | 61.51 | 0.99 | 0.96 |
|  | *Luetkea pectinata* | 1578.5 | 82.28 | 1.00 | 1.00 |
|  | *Juncus parryi* | 1644.96 | 67.76 | 0.98 | 0.97 |
|  | *Hieracium triste* | 1709.74 | 56.82 | 0.99 | 0.96 |
|  | *Pedicularis contorta* var. *contorta* | 1709.74 | 50 | 1.00 | 0.98 |
|  | *Penstemon procerus* var. *tolmiei* | 1853.38 | 61.59 | 0.98 | 0.97 |
|  | *Tsuga mertensiana* | 1853.38 | 56.67 | 1.00 | 0.97 |
|  | *Sedum divergens* | 1989.35 | 68.84 | 1.00 | 1.00 |
|  | *Campanula parryi* var. *idahoense* | 1989.35 | 64.29 | 1.00 | 1.00 |
|  | *Veronica cusickii* | 2203.53 | 70.62 | 1.00 | 0.99 |
|  | *Vaccinium cespitosum* | 2203.53 | 50 | 1.00 | 0.96 |
|  | *Montia parvifolia* | 2390.14 | 55.56 | 1.00 | 0.96 |
|  | *Lomatium martindalei* | 2653.88 | 86.67 | 1.00 | 1.00 |
|  | *Micranthes ferruginea* | 2653.88 | 76.29 | 1.00 | 0.98 |
| **Decreasers** | *Allium crenulatum* | 1644.96 | 46.67 | 1.00 | 0.97 |
|  | *Erigeron compositus* | 1853.38 | 71.43 | 1.00 | 1.00 |
|  | *Heuchera cylindrica* | 1853.38 | 71.93 | 1.00 | 1.00 |
|  | *Polemonium pulcherrimum* var. *pulcherrimum* | 1853.38 | 76.19 | 1.00 | 1.00 |
|  | *Packera streptanthifolia* | 1989.35 | 60.22 | 0.98 | 0.99 |
|  | *Androsace nivalis* | 1989.35 | 55.56 | 1.00 | 1.00 |
|  | *Silene suksdorfii* | 2458.29 | 68 | 0.98 | 0.96 |
|  | *Elymus elymoides* ssp. *brevifolius* | 2653.88 | 88.46 | 1.00 | 0.99 |
|  | *Castilleja elmeri* | 2653.88 | 73.08 | 0.98 | 0.97 |

## Discussion

We sought to elucidate patterns in alpine plant communities across the Cascade Mountain Range in Washington to link environmental variables to dissimilarity, compare results using taxonomic and phylogenetic dissimilarity, and identify species linked to significant patterns. We found significant spatial autocorrelation of alpine plant community composition (i.e., peaks geographically closer to each other and sampled in similar years/months were more likely to have similar plant communities). A suite of geologic, climatic, topographic, and spatial explanatory variables were identified as consistent predictors of composition, foremost of which was the age at which each peak’s geologic parent material was formed. Patterns were markedly consistent across both taxonomic and phylogenetic distance as well, despite phylogeny monotonically decreasing dissimilarity between peaks. A plethora of plant species were identified as indicators of these geologic and climatic patterns, revealing adaptational processes specific to the unique, harsh alpine ecosystems.

## Drivers of Dissimilarity

The existence of alpine plant communities on disconnected summit patches separated by a matrix of subalpine and montane forest have led researchers to view their beta-diversity in the context of island biogeography, where dispersal between summits increases dissimilarity at greater distances [13,14]. This relationship was apparent in the Cascades, with dissimilarity highly correlated to distance (Table 1), thus we can infer that dispersal likelihood decreases with distance at the mountain range level. These results of autocorrelation in alpine plant communities do not directly answer questions of determinism since environmental gradients are simultaneously spatially autocorrelated. Rather, our later investigation of environmental variables can be understood in the context of a dispersal-limited system, where peaks closer to one another have more alike plant communities.

The age of formation for each peak’s geologic parent material had the greatest influence on alpine plant community composition, accounting for 8-18% of the variation in our data (Tables 2 and 3). Washington’s alpine geology can be understood through the three geologic periods in which it was formed: diverse Paleozoic and Mesozoic bedrock formed undersea and transported as terranes create a matrix on which Cenozoic volcanoes have erupted new material. This mosaic of geologic diversity of the Cascade Mountain range was hypothesized to drive previous findings of nonconformity to environmental gradients compared to the Rocky Mountains and other North America mountain ranges [19,26] . Our results support this hypothesis and demonstrate the potential influence of geology on alpine plants at a mountain-range level that has not been often incorporated into biogeographical studies. Due to the conflation of variance and unquantified complexities that could predict alpine community composition [e.g., 23], determining the degree of stochasticity is inherently difficult. By including geology in our analyses, we clearly illustrated deterministic patterns enacted on a mountain-range level in the region.

An especially distinct geological group that emerged from our analyses was ultramafic parent material formed in the Paleozoic Era, which produces serpentine soils with uniquely high concentrations magnesium and other heavy metals that are toxic to most plants. Plants that do survive on serpentine soils experience high rates of endemism, with traits evolving independently multiple times within species and affecting traits that alter reproductive phenology and enhance drought-tolerance [48,49]. In our broad survey of alpine plants across the Cascade Range in Washington, the six Paleozoic and ultramafic peaks have the most distinct plant community, forming a well-separated and clustered group in our NMDS (Fig 5).

Out of the six climate variables we examined, three had significant influences on alpine plant community composition (Table 2, Fig 5). Mean annual precipitation had the strongest effect of the three, followed by the temperature of the wettest quarter (December, January, and February) and mean annual temperature. In combination, thee three variables represent observed climatic trends across the range manifest in its vegetation patterns. Peaks on the east side of the Cascade crest receive far less precipitation due to the orographic effect [25] and resultingly have more sparse, dryland-adapted species than wetter peaks in the west. In our study area, annual mean and winter temperatures are primarily a product of elevation, with higher-elevation peaks having colder temperatures and alpine-tundra vegetation [1]. Each of these three climatic variables are expected to shift with projected climate change in the Cascade Range [50]; our evidence is consistent with the literature in suggesting this could mediate drastic transitions of alpine plant communities [3,5,18].

With the strength of spatial, geologic, and climatic model terms, we conclude the alpine plant communities of the Cascade Mountain Range are deterministically formed despite their isolation. This insight is critical to management, informing site prioritization (for conservation or outplanting) with knowledge that geology and climate have and will shape the survival of alpine plants.

## Phylogenetic Dimensions

Inclusion of phylogenetic distances added a valuable dimension to our results, conferring evolutionary relationships determining today’s lineages. We found constantly lower phylogenetic distances than taxonomic (Fig 4), consistent with others’ conclusions of phylogenetic niche conservatism: closely related species are more likely to occur in similar environmental conditions [9]. For spatial autocorrelation, using phylogenetic distance translated to slightly lower effect strength and significance (Table 1), indicating a higher prevalence of environmental variables sorting plant communities at genus or family levels. This is somewhat expected as dispersal limitation is less relevant when not being limited by each species’ dispersal range, demonstrating the strength of comparing phylogenetic distance to answer biogeographic questions.

Contrasting to the results of Marx et al. (24), we found tree-wide phylogenetic structuring: maintained relationships of distance, elevation, geologic parent material, and precipitation variability with phylogenetic distance. These results suggest the influential patterns on alpine plants have created evolutionarily distinct lineages [22] with communities structured by the same spatial and environmental factors that originally drove their formation. This tracks with our Paleozoic plant group’s phenological reproductive isolation and frequent independent evolution [48] to create a unique community both evolutionarily and functionally. The evolutionary independence of this and other ecosystem demands special attention into the future as the impacts of human development (here, recreation) and climate change can imperil the small extent of this edaphic habitat. Potential further research is abundant in the linkages between phylogenies (or deeper genomics) within biogeographical patterns and in the context of conservation.

## Indicator Species

We used indicator species analysis to further explore the significant biogeographical influences on alpine plant community composition across our study area. This yielded valuable insights into community structure and underlying morphological traits. We identified 36 species indicating our geologic parent material periods of formation, almost half of which were associated with Paleozoic peaks. This Paleozoic species pool includes two interesting groups:serpentine-endemic or constrained species, and widespread or lowland species. Endemic species like *Aspidotis densa* (serpentine fern), *Polystichum lemmonii* (Shasta fern), and *Poa curtifolia* (Wenatchee bluegrass) are, of course, expected since their distribution within our study area is restricted to this region only and they possess adaptations to serpentine conditions. Lowland and widespread species like *Eriophyllum lanatum* (woolly sunflower) and *Pseudotsuga menziesii* (Douglas-fir) only occurring in these alpine areas are curious and may suggest serpentine treeline alteration or other convergent adaptations to exist both ubiquitously and to exist in harsh conditions.

We identified 23 species that indicate affinities to different levels of mean annual precipitation across our study area, 14 of which are associated with high precipitation and nine with low precipitation. Broadly, species on peaks high in precipitation belong to a subarctic heath vegetation communities characterized by species like *Vaccinium cespitosum* (dwarf huckleberry) and *Luetkea pectinata* (partridgefoot). These communities are higher in total biomass and have higher competition for space, potentially arising from more abundant water resources. Another subset of species within this pool are typical of mesic, mossy bedrock (e.g., *Micranthes ferruginea* and *Montia parvifolia*) that have high moisture requirements. Species on peaks low in precipitation more closely fit expected alpine plants’ adaptations to scarce resources, such as having taproots (e.g., *Eriogonum pyrolifolium*), succulent growth forms (e.g., *Packera streptanthifolia*) and underground storage structures (e.g., *Allium crenulatum*). These morphological variations across a water stress gradient serve to “ground-truth” our biogeographical findings and act as examples of how these broad patterns translate to plant adaptations. A fruitful research direction may be quantifying this morphological variation as functional traits in comparison to mountain-range level environmental gradients.

Spanning biogeographical, evolutionary, and morphological relationships, we found patterns in space, geology, and climate that shape the alpine plant communities of the Cascade Mountain Range. There is ample opportunity to continue this exploration with greater coverage of peaks in the Cascades and beyond, and with deeper methods of plant genomes, traits, and biotic/abiotic influences. Understanding these relationships is vital to both our understanding of ecosystem processes in the alpine and how best to conserve them.

## Supporting information

Full Reproducible Code

## Acknowledgements

Thank you to all past interns who have been instrumental to the extent and quality of these data: Inthone Chao, Sophia Ronan, Brielle Canares, Samantha Lieberman, Phoebe Smurthwaite, Leo Linder, and Ava Kloss-Schmidt. We thank Hannah Marx and Eric DeChaine for assistance with project design. We also thank the following individuals for support throughout the project, Gary Brill, Tony Cavalieri, Peter Dunwiddie, Jim Duemmel, Noel Knoke, Ellen Look, Carol Nygren, Berl and Karen Nussbaum, Richard G. Olmstead, Peg Pearson, Laura Potash, Richard Ramsden, Don and Ann Schaechtel, Peter Stekel and Jennie Goldberg, and Donovan and Linda Tracy, and Peter Zika.

---

**Table S1.**
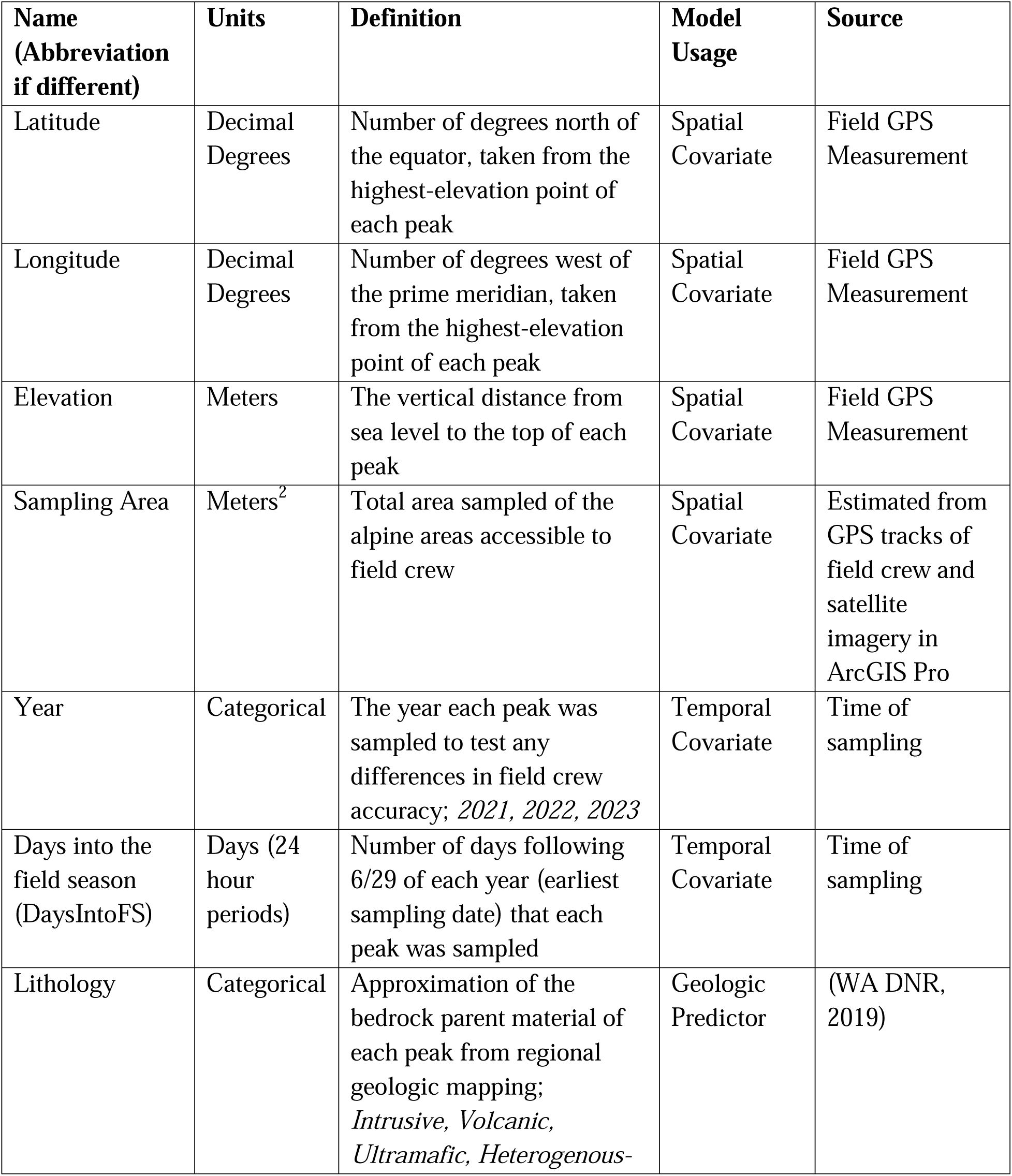
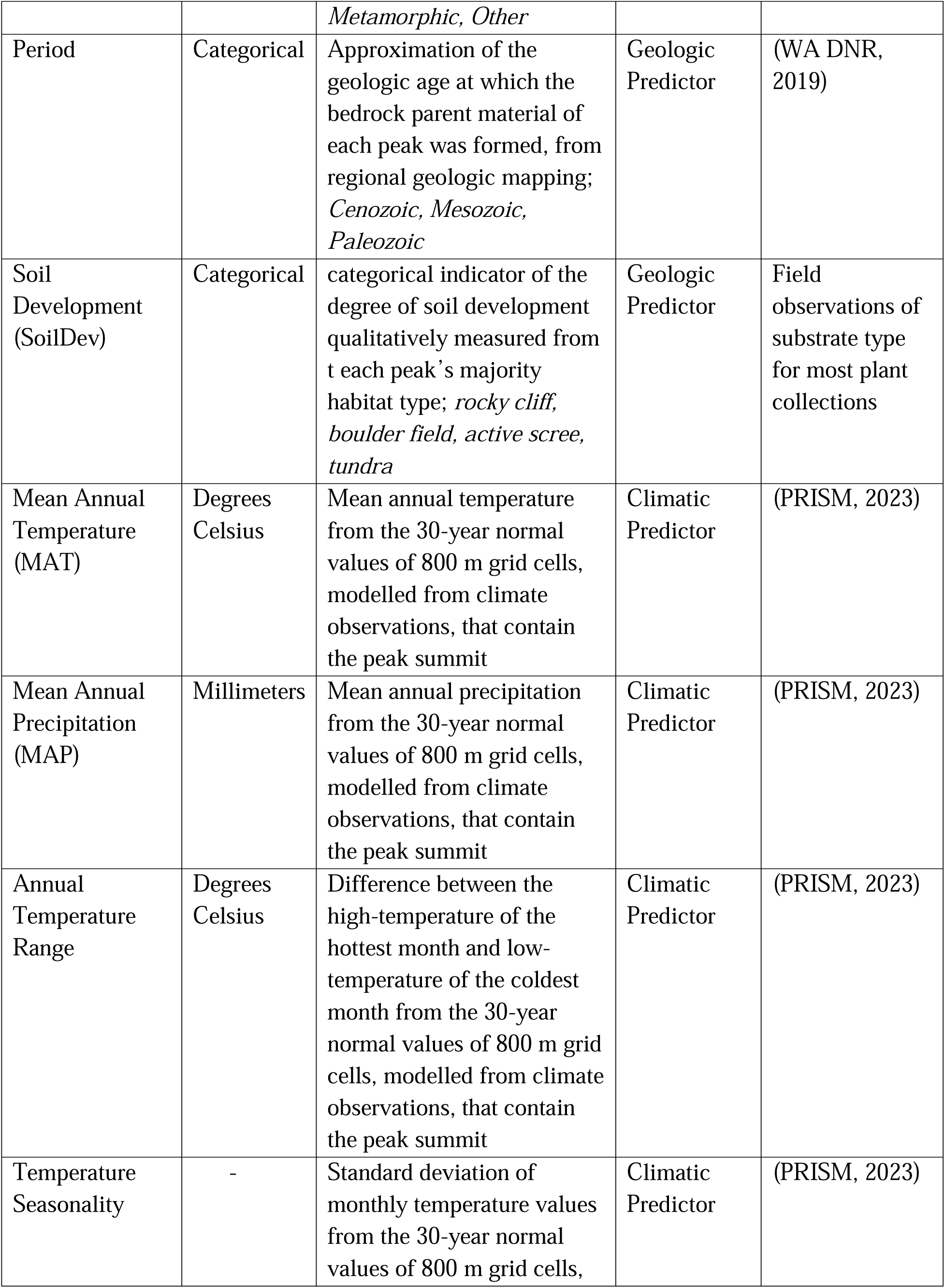
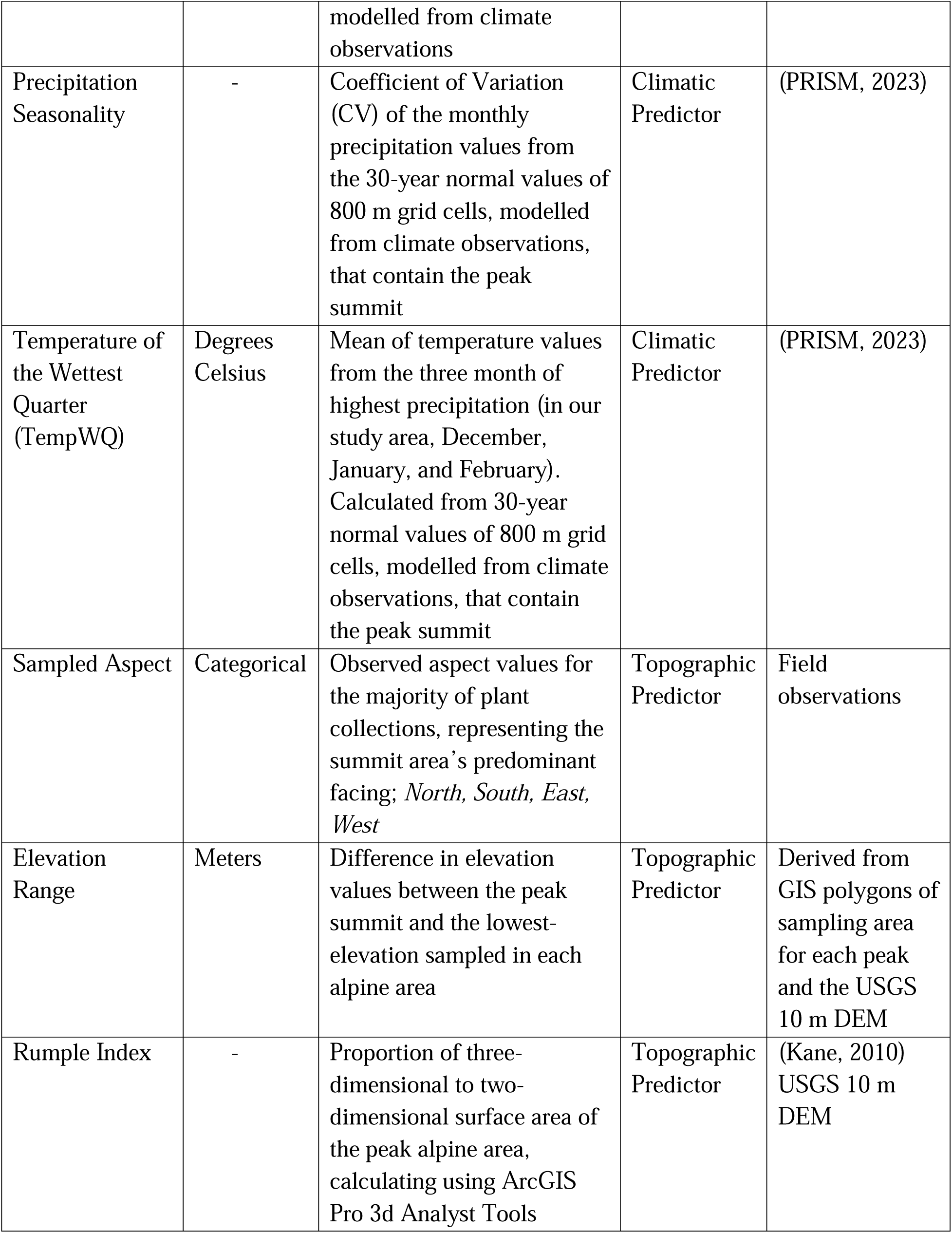
Model variables. Descriptions of all variables tested in PERMANOVA model analyses, including each’s unit, definition, source, and usage in the model selection framework.

