## Supplemental Table 1 for "Geology and climate drive alpine plant compositional variation among peaks in the Cascade Range of Washington"

**Supporting Information**

**Table S1. Model variables.** Descriptions of all variables tested in PERMANOVA model analyses, including each’s unit, definition, source, and usage in the model selection framework.

| **Name (Abbreviation if different)** | **Units** | **Definition** | **Model Usage** | **Source** |
| --- | --- | --- | --- | --- |
| Latitude | Decimal Degrees | Number of degrees north of the equator, taken from the highest-elevation point of each peak | Spatial Covariate | Field GPS Measurement |
| Longitude | Decimal Degrees | Number of degrees west of the prime meridian, taken from the highest-elevation point of each peak | Spatial Covariate | Field GPS Measurement |
| Elevation | Meters | The vertical distance from sea level to the top of each peak | Spatial Covariate | Field GPS Measurement |
| Sampling Area | Meters^2^ | Total area sampled of the alpine areas accessible to field crew | Spatial Covariate | Estimated from GPS tracks of field crew and satellite imagery in ArcGIS Pro |
| Year | Categorical | The year each peak was sampled to test any differences in field crew accuracy; *2021, 2022, 2023* | Temporal Covariate | Time of sampling |
| Days into the field season (DaysIntoFS) | Days (24 hour periods) | Number of days following 6/29 of each year (earliest sampling date) that each peak was sampled | Temporal Covariate | Time of sampling |
| Lithology | Categorical | Approximation of the bedrock parent material of each peak from regional geologic mapping; *Intrusive, Volcanic, Ultramafic, Heterogenous-Metamorphic, Other* | Geologic Predictor | (WA DNR, 2019) |
| Period | Categorical | Approximation of the geologic age at which the bedrock parent material of each peak was formed, from regional geologic mapping; *Cenozoic, Mesozoic, Paleozoic* | Geologic Predictor | (WA DNR, 2019) |
| Soil Development (SoilDev) | Categorical | categorical indicator of the degree of soil development qualitatively measured from t each peak’s majority habitat type; *rocky cliff, boulder field, active scree, tundra* | Geologic Predictor | Field observations of substrate type for most plant collections |
| Mean Annual Temperature (MAT) | Degrees Celsius | Mean annual temperature from the 30-year normal values of 800 m grid cells, modelled from climate observations, that contain the peak summit | Climatic Predictor | (PRISM, 2023) |
| Mean Annual Precipitation (MAP) | Millimeters | Mean annual precipitation from the 30-year normal values of 800 m grid cells, modelled from climate observations, that contain the peak summit | Climatic Predictor | (PRISM, 2023) |
| Annual Temperature Range | Degrees Celsius | Difference between the high-temperature of the hottest month and low-temperature of the coldest month from the 30-year normal values of 800 m grid cells, modelled from climate observations, that contain the peak summit | Climatic Predictor | (PRISM, 2023) |
| Temperature Seasonality |  | Standard deviation of monthly temperature values from the 30-year normal values of 800 m grid cells, modelled from climate observations | Climatic Predictor | (PRISM, 2023) |
| Precipitation Seasonality |  | Coefficient of Variation (CV) of the monthly precipitation values from the 30-year normal values of 800 m grid cells, modelled from climate observations, that contain the peak summit | Climatic Predictor | (PRISM, 2023) |
| Temperature of the Wettest Quarter (TempWQ) | Degrees Celsius | Mean of temperature values from the three month of highest precipitation (in our study area, December, January, and February). Calculated from 30-year normal values of 800 m grid cells, modelled from climate observations, that contain the peak summit | Climatic Predictor | (PRISM, 2023) |
| Sampled Aspect | Categorical | Observed aspect values for the majority of plant collections, representing the summit area’s predominant facing; *North, South, East, West* | Topographic Predictor | Field observations |
| Elevation Range | Meters | Difference in elevation values between the peak summit and the lowest-elevation sampled in each alpine area | Topographic Predictor | Derived from GIS polygons of sampling area for each peak and the USGS 10 m DEM |
| Rumple Index |  | Proportion of three-dimensional to two-dimensional surface area of the peak alpine area, calculating using ArcGIS Pro 3d Analyst Tools | Topographic Predictor | (Kane, 2010)  USGS 10 m DEM |
