## Supplementary material for "Geology and climate drive alpine plant compositional variation among peaks in the Cascade Range of Washington": Full Reproducible Code

**Full Reproducible Code (R version 4.4.2)**

```
### Geology and climate drive alpine plant compositional variation
### among peaks in the Cascade Mountain Range
## Erik W. Ertsgaard et al.
#Script prepared for publication in PLOS ONE

# 1.0 LOAD ITEMS -----------------------------------------------------------
# 1.1 Load packages ----

library(dplyr)
library(doParallel)
library(labdsv)
library(vegan)
library(tidyverse)
library(dismo)
library(phytools)
library(pez)
library(V.PhyloMaker2)
library(ggtree)
library(GUniFrac)
library(ggordiplots)
library(cowplot)
library(ggasym)
library(ggrepel)
library(indicspecies)
library(TITAN2)
library(ggpubr)

# 1.2 Load functions ----

# Custum, readable theme for figures
theme_custom <- theme_bw() +
  theme(axis.line = element_blank(),
        axis.ticks = element_blank(),
        axis.title = element_blank(),
        axis.text = element_blank())


# Pairwise comparisons for levels within PERMANOVA results, made by JDB
pairwise.adonis2 <- function(resp, fact, p.method = "none", nperm = 9999) {
  require(vegan)
  resp <- as.matrix(resp)
  fact <- factor(fact)
  fun.p <- function(i, j) {
    fact2 <- droplevels(fact[as.numeric(fact) %in% c(i, j)])
    index <- which(fact %in% levels(fact2))
    resp2 <- as.dist(resp[index, index])
    result <- adonis2(resp2 ~ fact2, permutations = nperm)
    result$`Pr(>F)`[1]
  }
  multcomp <- pairwise.table(fun.p, levels(fact), p.adjust.method = p.method)
  return(list(fact = levels(fact), p.value = multcomp, p.adjust.method = p.method))
}


# Model selection for phylogenetic distance matrix, built from fit_models base code in AICcPermanova package (https://rdrr.io/cran/AICcPermanova/src/R/fit_models.R)
fit_models_u <- function(all_forms,
                         unifrac_distance,
                         env_data,
                         ncores = 2,
                         log = TRUE,
                         logfile = "log.txt",
                         multiple = 100,
                         strata = NULL,
                         verbose = FALSE){
  AICc <- R2 <- term <- x <- NULL
  if(log){
    if(file.exists(logfile)){
      file.remove(logfile)
    }
  }

  meta_data <- all_forms
  if(!("max_vif" %in% colnames(meta_data))){
    meta_data$max_vif <- NA
  }

  # Check for missing values
  missing_rows <- !complete.cases(env_data)

  if (any(missing_rows)) {
    if(verbose){
      # Print message about missing rows and columns
      message(sprintf("Removing %d rows with missing values\n", sum(missing_rows)))
      message("Columns with missing values: ")
      message(names(env_data)[colSums(is.na(env_data)) > 0], sep = ", ")
    }
  }

  # Filter out missing rows
  new_env_data <- env_data[complete.cases(env_data), ]


  cl <- parallel::makeCluster(ncores)
  doParallel::registerDoParallel(cl)

  Distance <- unifrac_distance

  Fs <- foreach(x = 1:nrow(meta_data), .packages = c("vegan", "dplyr", "AICcPermanova", "tidyr", "broom", "doParallel"), .combine = bind_rows, .export = c("Distance")) %dopar% {

    Response = new_env_data
    Response$y <- rnorm(n = nrow(Response))
    gc()

    Temp <- meta_data[x,]

    if(is.null(strata)){
      Model <- try(vegan::adonis2(as.formula(Temp$form[1]), data = Response, by = "margin"))
    }

    if(!is.null(strata)){
      # Convert strata variable to factor
      strata_factor <- factor(Response[[strata]])
      Model <- try(with(Response, vegan::adonis2(as.formula(Temp$form[1]), data = Response, by = "margin", strata = strata_factor)), silent = TRUE)
    }


    Temp <- tryCatch(
      expr = cbind(Temp, AICcPermanova::AICc_permanova2(Model)),
      error = function(e) NA
    )


    if(is.na(Temp$max_vif)){
      Temp$max_vif <- tryCatch(
        expr = VIF(lm(as.formula(stringr::str_replace_all(Temp$form[1], "Distance ", "y")), data = Response)),
        error = function(e) NA
      )

    }

    Rs <- tryCatch(
      {
        tidy_model <- broom::tidy(Model)
        if (inherits(tidy_model, "try-error")) {
          stop("Error occurred in broom::tidy(Model)")
        }
        tidy_model |>
          dplyr::filter(!(term %in% c("Residual", "Total"))) |>
          dplyr::select(term, R2) |>
          tidyr::pivot_wider(names_from = term, values_from = R2)
      },
      error = function(e) {
        message("Error: ", conditionMessage(e))
        NULL
      }
    )


    if(log){
      if((x %% multiple) == 0){
        sink(logfile, append = TRUE)
        cat(paste("finished", x, "number of models", Sys.time(), "of",  nrow(meta_data)))
        cat("\n")
        sink()
      }

    }

    Temp <- bind_cols(Temp, Rs)
    Temp
  }
  parallel::stopCluster(cl)
  Fs <- Fs |>
    dplyr::arrange(AICc)
  return(Fs)
}


# 1.3 Load data ----

Alpine_Data <- read.csv("data/Cascades_Master_33x338.csv",
                        header = TRUE,
                        row.names = 1)

dim(Alpine_Data) #matches expected dimensionality

sp.list <- read.csv("data/Cascades_Code_Key_308x3.csv")

#Optimizing sp.list for phylo.maker()
colnames(sp.list) <- c("Taxon", "Code2Letter", "Family")
sp.list$genus <- word(sp.list$Taxon, 1)
sp.list <- sp.list[ ,c(1,4,3)]

#Changing classifications to fit with backbone tree used in phylo.maker()
sp.list[sp.list=="Antennaria lanata"] <- "Antennaria carpatica var. lanata"
sp.list[sp.list=="Erythranthe caespitosa"] <- "Erythranthe tilingii var. caespitosa"
sp.list[sp.list=="Polystichum lemmonii"] <- "Polystichum mohrioides var. lemmonii"
sp.list[sp.list=="Hydrophyllaceae"] <- "Boraginaceae" #Hydrophyllaceae is not recognized. This affects Phacelia and Romanzoffia.
sp.list[sp.list=="Valerianaceae"] <- "Caprifoliaceae" #Valerianaceae is not recognized. This affects Valeriana.


species_matrix <- Alpine_Data[ , 1:307] %>%
  replace(is.na(.), 0)

explanatory_matrix <- Alpine_Data[ ,308:337]


explanatory_matrix$TempSeasonality <- explanatory_matrix$TempSD*100

explanatory_matrix$PrecipSeasonality <- explanatory_matrix$PrecipSD/(explanatory_matrix$MAP/12)

explanatory_matrix <- explanatory_matrix %>%
  mutate(TempWQ = ((MinDecT + MinJanT + MinFebT + MaxDecT + MaxJanT + MaxFebT)/6))

# 1.4 Data Adjustments ----

explanatory_matrix_rel <- explanatory_matrix %>%
  dplyr::select(-c(SampledAspect, Lithology, Period, RockCode, SoilDev, DateSampled, CrestPosition, Year)) %>%
  decostand(method = "standardize", MARGIN = 2) %>% #relativizing continuous predictors to Z-scores
  cbind(explanatory_matrix[ , c("SampledAspect", "Lithology", "Period", "RockCode", "SoilDev", "DateSampled",
                                "CrestPosition", "Year")])


explanatory_matrix_rel %>%
  dplyr::select(Latitude, Longitude, Elevation, DaysIntoFS, SamplingArea, MAT, MAP, AnnTempRange,
                TempWQ, TempSeasonality, PrecipSeasonality, ElevationRange, Rumple) %>%
  cor() %>%
  as.data.frame() %>%
  write.csv(file = "Correlation_Matrix.csv")

explanatory_matrix_rel$Year <- as.factor(explanatory_matrix_rel$Year)


# 2.0 Phylogenetic Tree  ----

# making tree
alpine.tre <- phylo.maker(sp.list,
                          tree = GBOTB.extended.WP,
                          nodes = nodes.info.1.WP,
                          scenarios = "S3")


alpine.tre$scenario.3$tip.label <- gsub("_", " ", alpine.tre$scenario.3$tip.label)

alpine.tre$scenario.3$tip.label[alpine.tre$scenario.3$tip.label=="Antennaria carpatica var. lanata"] <- "Antennaria lanata"
alpine.tre$scenario.3$tip.label[alpine.tre$scenario.3$tip.label=="Erythranthe tilingii var. caespitosa"] <- "Erythranthe caespitosa"
alpine.tre$scenario.3$tip.label[alpine.tre$scenario.3$tip.label=="Polystichum mohrioides var. lemmonii"] <- "Polystichum lemmonii"


write.tree(alpine.tre$scenario.3, "speciesturnover.tre")


# Unifrac distance
comm.wide <- as.matrix(species_matrix)

colnames(comm.wide) <- alpine.tre$scenario.3$tip.label

unifrac_distance <- GUniFrac(comm.wide, alpine.tre$scenario.3, alpha = c(0, 0.5, 1)) %>%
  .$unifracs %>%
  .[, , "d_UW"]

write.csv(unifrac_distance, file = "Unweighted UniFrac Distances.csv")


# 3.0 Mantel Test -------------------------

geog_distance <- explanatory_matrix[, c("Latitude", "Longitude")] %>%
  vegdist(method = "euclidean")

# 3.1 Mantel - Jaccard ----
jaccard_distance <- species_matrix %>%
  vegdist(method = "jaccard")

set.seed(13)
J.mantel <- mantel(xdis = jaccard_distance,
                   ydis = geog_distance,
                   method = "pearson",
                   permutations = 99999)
J.mantel

# 3.2 Mantel - Unifrac ----
set.seed(13)
U.mantel <- mantel(xdis = unifrac_distance,
                   ydis = geog_distance,
                   method = "pearson",
                   permutations = 99999)
U.mantel

# 4.0 PERMANOVA --------------------

# 4.1 PERMANOVA & AICc Model Selection - Jaccard -----

##JDB - start by testing each variable separately and only retaining those that are significant
explan <- c("Latitude", "Longitude", "Elevation", "SamplingArea",
            "Year", "DaysIntoFS",
            "Lithology", "Period", "SoilDev",
            "SampledAspect", "ElevationRange", "Rumple",
            "MAT", "MAP", "AnnTempRange", "TempSeasonality", "PrecipSeasonality", "TempWQ")
res <- c()
for(i in 1:length(explan)) {
  formula <- as.formula(paste("species_matrix ~ ", explan[i]))
  set.seed(42)
  model <- adonis2(formula = formula,
                   data = explanatory_matrix_rel,
                   distance = species_matrix,
                   method = "jaccard")
  res.temp <- data.frame(variable = explan[i],
                         R2 = model$R2[1],
                         P_value = model$`Pr(>F)`[1])
  res <- rbind(res, res.temp)
}

## SampledAspect and ElevationRange not significant when tested individually
explan <- explan[! explan %in% c("SampledAspect", "ElevationRange")]

##JDB - fit rest of variables
models <- make_models(vars = explan,
                      ncores = 2) %>%
  filter_vif(env_data = explanatory_matrix_rel)

set.seed(42)
fitted.models <- fit_models(all_forms = models,
                            veg_data = species_matrix,
                            env_data = explanatory_matrix_rel,
                            ncores = 4,
                            method = "jaccard")
## took 2-3 hours to run on my laptop

final.models <- select_models(fitted.models, delta_aicc = 2) #184 of the 65536 possible models

AICc.J <- akaike_adjusted_rsq(final.models) %>%
  dplyr::filter(!is.na(AICc.J$Full_Akaike_Adjusted_RSq)) #Period, MAP, Latitude, TempWQ, MAT, Rumple


#Pairwise lithology
set.seed(13)
p.a.j <- pairwise.adonis2(jaccard_distance, explanatory_matrix$Lithology)

write.csv(as.data.frame(p.a.j$p.value), file = "Pairwise_Adonis_Jaccard.csv")

# 4.2 PERMANOVA & AICc Model Selection - UniFrac -----

##JDB - start by testing each variable separately and only retaining those that are significant
res2 <- c()
for(i in 1:length(explan)) {
  formula <- as.formula(paste("unifrac_distance ~ ", explan[i]))
  set.seed(42)
  model <- adonis2(formula = formula,
                   data = explanatory_matrix_rel,
  )
  res.temp <- data.frame(variable = explan[i],
                         R2 = model$R2[1],
                         P_value = model$`Pr(>F)`[1])
  res2 <- rbind(res, res.temp)

}

## SampledAspect and ElevationRange not significant when tested individually, can use same variable set as Jaccard


# using customized fit_models function compatible with unifrac distances to fit variables as above
set.seed(42)
fitted.models.2 <- fit_models_u(all_forms = models,
                                unifrac_distance = unifrac_distance,
                                env_data = explanatory_matrix_rel,
                                ncores = 4)

# selectiong models with deltaAICc values less than 2
final.models.2 <- select_models(fitted.models.2, delta_aicc = 2) #58 of the 65536 possible models

AICc.U <- akaike_adjusted_rsq(final.models.2) %>%
  dplyr::filter(!is.na(AICc.U$Full_Akaike_Adjusted_RSq)) # same final variables as jaccard, Period more predictive


# making table for final model-averaged results (Table 2)
data.frame(AICc.J$Variable, AICc.J$Full_Akaike_Adjusted_RSq, AICc.U$Full_Akaike_Adjusted_RSq) %>%
  write.csv("AICc_results_table.csv")


# 5.0 NMDS ------------------------

# 5.1 NMDS - Jaccard ----
## Jaccard
set.seed(13)
jaccard.NMDS <- metaMDS(comm = species_matrix,
                        autotransform = FALSE,
                        distance = "jaccard",
                        engine = "monoMDS",
                        k = 3,
                        model = "global",
                        maxit = 400,
                        try = 40,
                        trymax = 100)

# 5.2 NMDS - Unifrac ----
unifrac.NMDS <- metaMDS(comm = unifrac_distance,
                        autotransform = FALSE,
                        engine = "monoMDS",
                        k = 3,
                        model = "global",
                        maxit = 400,
                        try = 40,
                        trymax = 100)

u.points <- data.frame(unifrac.NMDS$points)


# 6.0 ISA ----
set.seed(13)
lith.ISA <- multipatt(x = species_matrix,
                      cluster = explanatory_matrix$Lithology,
                      duleg = FALSE,
                      permutations = 9999,
)
summary(lith.ISA,
        minstat = 0.7072)

# 7.0 TITAN ----
species_filtered <- species_matrix %>%
  vegtab(minval = 3) #removing species found on less than 3 peaks

set.seed(13)
preseas.TITAN <- titan(env = explanatory_matrix_rel$PrecipSeasonality,
                       txa = species_filtered,
                       numPerm = 1000,
                       nBoot = 1000)

preseas.TITAN$sppmax %>%
  as.data.frame() %>%
  dplyr::select(zenv.cp, freq, maxgrp, IndVal, purity, reliability, filter) %>%
  filter(filter != 0) %>%
  arrange(maxgrp, zenv.cp)

# 8.0 Graphics ----
# 8.1 Phylogeny Figures ----
## phylogenetic tree
# Legend
majorclades<- data.frame(node = c(536, 545, 526, 494, 312),
                         Classification = c(" ", " ", "   ", "    ", "     "))

#Orange = "Non-seed plants"
#Green = "Gymnosperms"
#Blue = "Monocots"
#Purple = "Eudicots"

clade.nodelabels <- c(rep("Eudicot", 225), rep("Monocot", 56), rep("Gymnosperm", 11), rep("Non-seed Plant", 15), rep(NA, 5), rep("Eudicot", 181), NA, rep("Monocot", 31), NA, rep("Gymnosperm", 9), NA, rep("Non-seed Plant", 8), NA, "Non-seed Plant")

plottree <- as_tibble(alpine.tre$scenario.3) %>%
  mutate(clade = clade.nodelabels) %>%
  as.treedata()

png(filename = "alpinetree.png", pointsize = 3, res = 700, width = 5000, height = 5000)
p <- ggtree(alpine.tre$scenario.3,
            ladderize=TRUE,
            layout = "circular",
            branch.length = "none",
            aes(color = plottree@data$clade)) +

  scale_color_manual(values = c('#e7298a','#d95f02', '#7570b3','#1b9e77' )) +

  geom_tiplab(size= 1, color = "black", hjust = -0.1, offset = 0.1) +

  geom_cladelab(node=319, label="Asteraceae", offset = 8, offset.text = 1, angle = "auto", fontsize = 2)+
  geom_cladelab(node=497, label="Poaceae", offset = 8, offset.text = 1, angle = "auto", fontsize = 2)+
  geom_cladelab(node=451, label="Brassicaceae", offset = 6, offset.text = 1, angle = "auto", fontsize = 2)+
  geom_cladelab(node=514, label="Cyperaceae", offset = 8.5, offset.text = 1, angle = "auto", fontsize = 2)+
  geom_cladelab(node=474, label="Saxifragaceae", offset = 8.5, offset.text = 1, angle = "auto", fontsize = 2)+
  geom_cladelab(node=381, label="Ericaceae", offset = 9.5, offset.text = 1, angle = "auto", fontsize = 2)+
  geom_cladelab(node=432, label="Rosaceae", offset = 9.5, offset.text = 1, angle = "auto", fontsize = 2)+

  theme(legend.position = c(0.93, 0.93))

p

dev.off()

# 8.2 Heatmap Figure ----
# unifrac
heatmapuni <- data.frame(unifrac_distance) %>%
  rownames_to_column() %>%
  gather(colname, value, -rowname)

u.hm <- ggplot(heatmapuni, aes(x = rowname, y = colname, fill = value)) +
  geom_tile() +
  theme(axis.text.x = element_text(angle = 90, vjust = 0.5, hjust=1), plot.title = element_text(hjust = 0.5)) +
  scale_y_discrete(position = "right") +
  labs(title = "Phylogenetic Distance", x = NULL, y = NULL) +
  scale_fill_viridis_c(limits = c(0,1), name = "Dissimilarity")
u.hm

# jaccard
heatmapjacc <- species_matrix %>%
  vegdist(method = "jaccard",
          upper = TRUE,
          diag = TRUE) %>%
  as.matrix(nrow = 32) %>%
  as.data.frame() %>%
  rownames_to_column() %>%
  gather(colname, value, -rowname)

j.hm <- ggplot(heatmapjacc, aes(x = rowname, y = colname, fill = value)) +
  geom_tile() +
  theme(axis.text.x = element_text(angle = 90, vjust = 0.5, hjust=1), plot.title = element_text(hjust = 0.5)) +
  labs(title = "Taxonomic Distance", x = NULL, y = NULL) +
  scale_fill_viridis_c(limits = c(0,1), name = "Dissimilarity")
j.hm

comparison.plot <- data.frame(heatmapjacc$value, heatmapuni$value)
c.plot <- ggplot(data = comparison.plot, aes(x = heatmapjacc.value, y = heatmapuni.value)) +
  geom_point(aes(fill = heatmapuni.value), size = 2.5, pch = 21) +
  scale_fill_distiller(palette = 16, name = "Dissimilarity") +
  xlim(0,1) + ylim(0,1) +
  labs(x = "Phylogenetic Distance", y = "Taxonomic Distance") +
  geom_abline(slope = 1, intercept = 0) +
  theme_bw()

hm.data <- data.frame(heatmapjacc, heatmapuni)
hm.data$value.diag <- rep(0, 32)


# Set the levels of colname and rowname to match the original dataframe order
hm.data$colname <- factor(hm.data$colname, levels = unique(hm.data$colname))
hm.data$rowname <- factor(hm.data$rowname, levels = unique(hm.data$rowname))

# Reorder the levels of colname and rowname according to the geology variable
hm.data$colname <- factor(hm.data$colname, levels = unique(hm.data$colname)[order(explanatory_matrix$Lithology)])
hm.data$rowname <- factor(hm.data$rowname, levels = unique(hm.data$rowname)[order(explanatory_matrix$Lithology)])


hm <- ggplot(hm.data, aes(x = colname, y = rowname, fill = value)) +
  geom_asymmat(aes(fill_tl = value, fill_br = value.1, fill_diag = value.diag)) +
  scale_fill_tl_distiller(palette = 16, limits = c(0, 1), name = "Dissimilarity") +
  scale_fill_br_distiller(palette = 16, limits = c(0, 1), name = "Dissimilarity") +
  scale_fill_diag_gradient(low = "#225ea8", high = "#225ea8") +
  guides(fill_diag = "none") +
  theme(axis.text.x = element_text(angle = 90, vjust = 0.5, hjust = 1),
        plot.title = element_text(hjust = 0.5),
        legend.direction = "horizontal",
        legend.title = element_text("none")) +
  labs(x = NULL, y = NULL)


# 8.3 NMDS Figures ----

##Jaccard
j.points <- data.frame(jaccard.NMDS$points)

# Period
j.l.nmds <- ggplot(j.points, aes(x = MDS1, y = MDS2,
                                 color = explanatory_matrix$Period,
                                 fill = explanatory_matrix$Period,
                                 shape = as.factor(explanatory_matrix$Lithology))) +
  stat_chull(mapping = aes(x = MDS1, y = MDS2,
                           color = explanatory_matrix$Period,
                           fill = explanatory_matrix$Period),
             geom = "polygon", alpha = 0.2, linewidth = 0.7,
             inherit.aes = FALSE) +
  scale_color_brewer(palette = "Set2") +
  scale_fill_brewer(palette = "Set2") +
  geom_point(size = 4) +
  guides(shape = guide_legend(title = "Type of Parent Material"),
         color = guide_legend(title = "Age of Parent Material Formation"),
         fill = "none") +
  labs(x = "NMDS1",
       y = "NMDS2") +
  theme_custom


# MAP
j.p.nmds <- ggplot(j.points, aes(x = MDS1, y = MDS2,
                                 fill = explanatory_matrix$MAP,
                                 size = explanatory_matrix$MAT)) +
  geom_point(pch = 21) +
  scale_fill_gradient(low = "khaki", high = "darkgreen") +
  guides(fill = guide_legend(title = "Mean Annual Precipitation (mm)"),
         size = guide_legend(title = expression(paste("Mean Annual Temperature (",degree~C,")")))) +
  labs(x = "NMDS1",
       y = "NMDS2") +
  theme_custom

pg1 <- plot_grid(j.g.nmds, j.l.nmds + theme(legend.position = "none"), j.p.nmds + theme(legend.position = "none"),
                 ncol = 3)

## Unifrac

# Period
u.l.nmds <- ggplot(u.points, aes(x = MDS1, y = MDS2,
                                 color = explanatory_matrix$Period,
                                 fill = explanatory_matrix$Period,
                                 shape = as.factor(explanatory_matrix$Lithology))) +
  stat_chull(mapping = aes(x = MDS1, y = MDS2,
                           color = explanatory_matrix$Period,
                           fill = explanatory_matrix$Period),
             geom = "polygon", alpha = 0.2, linewidth = 0.7,
             inherit.aes = FALSE) +
  scale_color_brewer(palette = "Set2") +
  scale_fill_brewer(palette = "Set2") +
  geom_point(size = 4) +
  guides(shape = guide_legend(title = "Type of Parent Material"),
         color = guide_legend(title = "Age of Parent Material Formation"),
         fill = "none") +
  labs(x = "NMDS1",
       y = "NMDS2") +
  theme_custom

# MAP
u.p.nmds <- ggplot(u.points, aes(x = MDS1, y = MDS2,
                                 fill = explanatory_matrix$MAP,
                                 size = explanatory_matrix$MAT)) +
  geom_point(pch = 21) +
  scale_fill_gradient(low = "khaki", high = "darkgreen") +
  guides(fill = guide_legend(title = "Mean Annual Precipitation (mm)"),
         size = guide_legend(title = expression(paste("Mean Annual Temperature (",degree~C,")")))) +
  labs(x = "NMDS1",
       y = "NMDS2") +
  theme_custom

# combined plots

plot(get_legend(j.l.nmds))
plot(get_legend(j.p.nmds))

pg3 <- plot_grid(j.l.nmds + theme(legend.position = "none"), j.p.nmds + theme(legend.position = "none"),
                u.l.nmds + theme(legend.position = "none"), u.p.nmds + theme(legend.position = "none"),
                 nrow = 2, byrow = FALSE)
pg3
```
